# Collecting reward to defend homeostasis: A homeostatic reinforcement learning theory

**DOI:** 10.1101/005140

**Authors:** Mehdi Keramati, Boris Gutkin

## Abstract

Efficient regulation of internal homeostasis and defending it against perturbations requires complex behavioral strategies. However, the computational principles mediating brain’s homeostatic regulation of reward and associative learning remain undefined. Here we use a definition of primary rewards, as outcomes fulfilling physiological needs, to build a normative theory showing how learning motivated behavior is modulated by the internal state of the animal. The theory proves that seeking rewards is equivalent to the fundamental objective of physiological stability, defining the notion of physiological rationality of behavior. We further give a formal basis for temporal discounting of reward. It also explains how animals learn to act predictively to preclude prospective homeostatic challenges, and attributes a normative computational role to the modulation of midbrain dopaminergic activity by hypothalamic signals.

## Introduction

Survival requires living organisms to maintain their physiological integrity within the environment. In other words, they must preserve homeostasis (e.g. body temperature, glucose level, etc.). Yet, how might an animal learn to structure its behavioral strategies to obtain the outcomes necessary to fulfill and even preclude homeostatic challenges? In this sense, efficient behavioral decisions depend on two brain circuits working in concert: the hypothalamic homeostatic regulation (HR) system, and the cortico-basal ganglia reinforcement learning (RL) mechanism. However, the computational mechanisms underlying this obvious coupling remain poorly understood, as the two systems have traditionally been studied separately.

On the one hand, classical negative feedback models of HR explain hypothalamic function in behavioral sensitivity to the “internal” state, by axiomatizing that animals minimize the deviation of some key physiological variables from their hypothetical setpoints (Marieb & Hoehn, 2012). To this end, a corrective response is triggered when a deviation from setpoint is sensed or anticipated (Sibly & McFarland, 1974; Sterling, 2012). A key lacuna in these models is how a simple corrective response (e.g. “go eat”) in response to a homeostatic deficit should be translated into a complex behavioral strategy for interacting with the dynamic and uncertain external world.

On the other hand, the computational theory of RL successfully explains the role of the cortico-basal ganglia system in behavioral adaption to the “external” environment, by exploiting experienced environmental contingencies and reward history (Rangel, Camerer, & Montague, 2008; Sutton & Barto, 1998). Critically, this theory is built upon one major axiom, namely, that the objective of behavior is to maximize reward acquisition. Yet, this suite of theoretical models does not resolve how the brain constructs the reward itself, and how the variability of the internal state impacts overt behavior.

Neurobiologically, accumulating evidence indicates intricate intercommunication between the hypothalamus and the reward-learning circuitry (Palmiter, 2007; Yeo & Heisler, 2012). The integration of the two systems is also behaviorally manifest in the classical behavioral pattern of anticipatory responding, in which animals learn to act predictively to preclude prospective homeostatic challenges. Here, we suggest an answer to the question of what computations, at an algorithmic level, are being performed in this biological integration of the two systems. Behaviorally, the theory explains anticipatory responding, extinction burst, and the rise-fall pattern of the response rate, as three behavioral phenomena for which the interaction between the two systems is necessary; thus, neither classical RL nor classical HR theories can account for these phenomena.

## Results

### Reward as need fulfillment

The term “reward” (equivalently: reinforcer, utility) has been at the heart of behavioral psychology since its foundation. In behavioral terms, reward refers to a stimulus that strengthens a desired response. According to RL theory, given the rewarding value of each outcome, animals learn the value of alternative choices as they experience them and receive feedback (i.e., reward) from the environment. However, animal behavior is variable even under well-controlled external conditions, suggesting that outcomes depend on the internal state.

The interaction between drive (as an internal state) and reward has been the subject of considerable debate in the motivation literature in psychology. Neo-behaviorists like Hull (Hull, 1943), Spence (Spence, 1956) and Mowrer (Mowrer, 1960) proposed the “drive-reduction” theory of motivation to define the nature of reward. According to this theory, one primary mechanism underlying reward is the usefulness of the corresponding outcome in fulfilling the homeostatic needs of the organism (Cabanac, 1971). Inspired by this theory, we derive a formal definition of primary reward as the ability of an outcome to restore the internal equilibrium of the physiological state. In the following sections, we demonstrate that our formal elaboration of the drive-reduction theory alleviates the criticisms raised against it.

We first define “homeostatic space” as a multidimensional metric space in which each dimension represents one physiologically regulated variable (the horizontal plane in Figure 1). The physiological state of the animal at each time *t* can be represented as a point in this space, denoted by *H_t_* = (*h*_1,_*_t_*, *h*_2,_*_t_*,.., *h_N,t_*), where *h_i,t_* indicates the state of the *i*-th physiological variable. For example, *h_i,t_* can refer to the animal’s glucose level, body temperature, plasma osmolality, etc. The homeostatic setpoint, as the ideal internal state, can be denoted as *H*\* = (*h*_1_^*^, *h*_2_^*^,.., *h_N_*^*^). As a mapping from the physiological to the motivational state, we define the “drive” as the distance of the internal state from the setpoint (the three-dimensional surface in Figure 1):

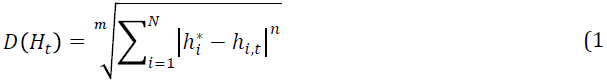

**Figure 1.**
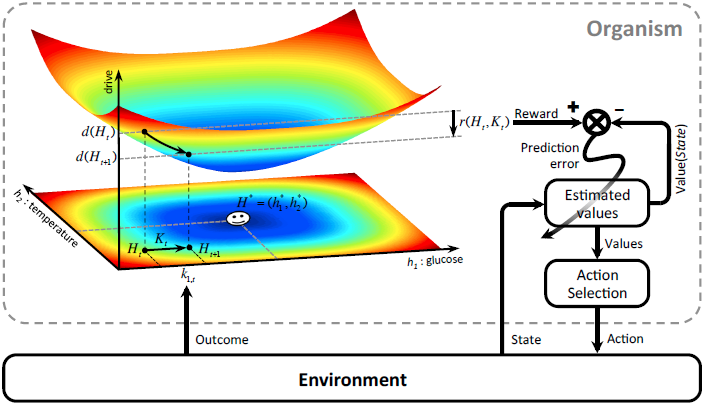
Schematics of the model in an exemplary two-dimensional homeostatic space. Upon performing an action, the animal receives an outcome *K_t_* from the environment. The rewarding value of this outcome depends on its ability to make the internal state, *H_t_*, closer to the homeostatic setpoint, *H*\*, and thus reduce the drive level (the vertical axis). This experienced reward, denoted by *r*(*H_t_, K_t_*), is then learned by a RL algorithm. Here a model-free RL algorithm (Daw, Niv, & Dayan, 2005) is shown in which a reward prediction error signal is computed by comparing the realized reward and the expected rewarding value of the performed response. This signal is then used to update the subjective value attributed to the corresponding response. Subjective values of alternative choices bias the action selection process.

Having defined drive, we can now provide a formal definition for primary reward based on drive reduction theory. Assume that as the result of an action, the animal receives an outcome *o_t_* at time *t*. The impact of this outcome on different dimensions of the animal’s internal state can be denoted by *K_t_* = (*k*_1,_*_t_*, *k*_2,_*_t_*,.., *k_N,t_*). For example, *k_i,t_* can be the quantity of glucose received as a result of outcome *o_t_*. Hence, the outcome results in a transition of the physiological state from *H_t_* to *H_t_*_+1_ = *H_t_* + *K_t_* (see Figure 1) and thus, a transition of the drive state from *D*(*H_t_*) to *D*(*H_t_*_+1_) = *D*(*H_t_* + *K_t_*). Accordingly, the rewarding value of this outcome can be defined as the consequent reduction of drive:

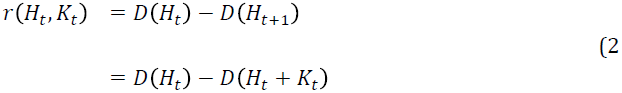

Intuitively, the rewarding value of an outcome depends on the ability of its constituting elements to reduce the homeostatic distance from the setpoint. We propose that this definition of reward is used by the brain’s reward learning machinery to structure behavior. Incorporating the physiological reward definition (Eq. 2) in a normative RL theory allows us to derive the major result of our theory, which is that the rationality of behavioral patterns is geared toward maintaining physiological stability.

### Rationality of the theory

Our definition of reward reconciles the RL and HR theories in terms of their normative assumptions: reward acquisition and physiological stability are mathematically equivalent behavioral objectives (see Materials and methods for the proof). More precisely, given the proposed definition of reward and given that animals discount future rewards (Chung & Herrnstein, 1967), any behavioral policy, *π*, that maximizes the sum of discounted rewards (*SDR*) also minimizes the sum of discounted deviations (*SDD*) from the setpoint, and vice versa. This can be represented as follows:

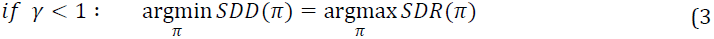

where *γ* is the discount factor. In this respect, reward acquisition sought by the RL system is an efficient means to guide an animal’s behavior toward satisfying the basic objective of defending homeostasis. Thus, our theory suggests a physiological basis for the rationality of reward seeking.

In the domain of animal behavior, one fundamental question is why animals should discount rewards the further they are in the future. Our theory indicates that reward seeking without discounting (*γ* = 1) would not lead, and may even be detrimental, to physiological stability (see Materials and methods). More precisely, in the absence of discounting, the rewarding value of behavioral policies that change the internal state only depends on the initial and final internal states, regardless of its trajectory in the homeostatic space. Thus, when *γ* = 1, the values of any two behavioral policies with equal net shifts of the internal state are equal, even if one policy moves the internal state along the shortest path, whereas the other policy results in large deviations of the internal state from the setpoint and threatens survival. These results hold for any form of temporal discounting (e.g., exponential, hyperbolic). In this respect, for the first time to our knowledge, our theory provides a normative explanation for the necessity of temporal discounting of reward: to maintain internal stability, it is necessary to discount future rewards.

### Anticipatory responding

A paradigmatic example of behaviors governed by the internal state is the anticipatory responses geared to preclude perturbations in regulated variables even before any physiological depletion (negative feedback) is detectable. Anticipatory eating and drinking that occur before any discernible homeostatic deviation (S C Woods & Seeley, 2002), anticipatory shivering in response to a cue that predicts the cold (Hjeresen, Reed, & Woods, 1986; Mansfield, Benedict, & Woods, 1983), and insulin secretion prior to meal initiation (S C Woods, 1991), are only a few examples of anticipatory responding.

One clear example of a conditioned homeostatic response is animals’ progressive tolerance to ethanol-induced hypothermia. Experiments suggest that this tolerance is mediated by associative learning processes (Mansfield & Cunningham, 1980): temperature deviations caused by ethanol injections decreased over trials, when injections preceded (i.e., were predictable) by a distinctive cue. Interestingly, in extinction trials where the ethanol was omitted, the animal temperature exhibited a significant increase above normal on cue presentation (Figure 2 - figure supplement 1). This result indicates that this tolerance is a conditioned response

Conditioned tolerance to the alcohol-induced homeostatic challenge clearly demonstrates that whether increasing the body temperature at any given time is rewarding or punishing depends on the internal state dynamics. Thus, the lack of internal state modulation of the rewarding value of responses in the classical RL theory renders it unable to explain anticipatory responding. Here we demonstrate that the integration of HR and RL processes can account for this phenomenon.

The model results (Figure 2) show that if a tolerance response (preventive increase in body temperature) precedes an ethanol injection, it results in a smaller deviation of body temperature from the setpoint, compared to the absence of the tolerance response (compare panels e and f of Figure 2). Therefore, expressing the tolerance response is the optimal behavior in terms of minimizing homeostatic deviation and thus, maximizing reward. As a result, the model gradually learns to choose this optimal strategy upon observing the cue (Figure 2b,c). In other words, the model learns to predictively generate a tolerance response, triggered by the conditioned stimulus, to minimize temperature deviation (see Materials and methods and Table S1 for simulation details). Thus, the model explains that the optimal homeostatic maintenance policy is acquired by associative learning mechanisms.

**Figure 2.**
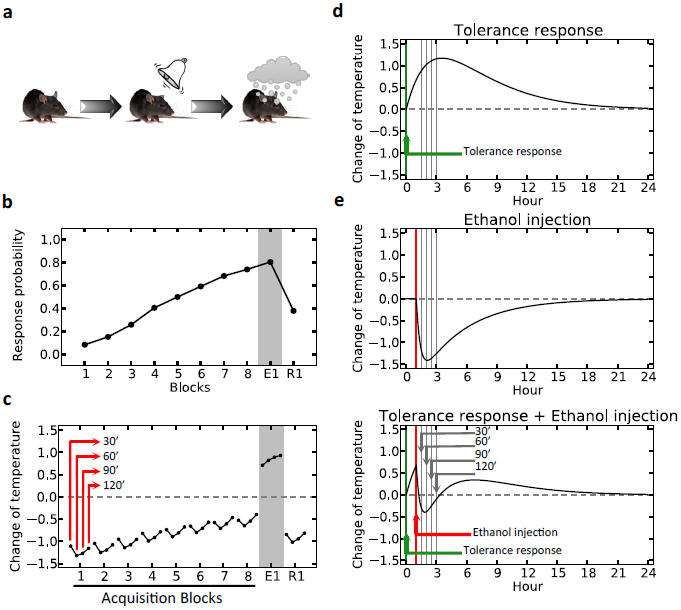
Anticipatory responding explained by the model. (a) In each block (day), the simulated agent receives an ethanol injection after the presentation of the stimulus. Plots (d) and (e) show the assumed dynamics of body temperature upon the initiation of the tolerance response and ethanol administration, respectively. Plot (f) shows the combined effect. Plot (b), which is averaged over 500 simulated agents, illustrates that the model gradually learns to choose the tolerance response after observing the stimulus. If the stimulus is not followed by ethanol injection, as in the first day of extinction (E1), it still triggers the tolerance response. However, the tolerance response is weakened after several extinction sessions, resulting in low response probability in the first day of re-acquisition (R1), where presentation of the cue is again followed by ethanol injection. (c) As in the experiment, the change in the body temperature is measured in every block, 30, 60, 90, and 120 min after ethanol administration. Plot (c) replicates experimental results (Figure S3b).

Our theory implies that animals are capable of learning not only Pavlovian learning (e.g. shivering, or tolerance to ethanol), but also instrumental anticipatory responding (e.g., pressing a lever to receive warmth, in response to a cold-predicting cue. See Figure S3). This is in contrast to the theory of predictive homeostasis where anticipatory behaviors are only *reflexive* responses to the predicted internal state upon observing cues (Sterling, 2012; Stephen C Woods & Ramsay, 2007).

### Extinction burst

Extinction is a procedure where the reinforcer maintaining a behavior is no longer provided upon performing that behavior. Although extinction eventually results in decreased response rate, a transient increase in the response rate is often observed in the early stages. This classical phenomenon, known as “extinction burst”, is demonstrated for a variety of reinforcers, in both animals and humans (Skinner, 1938). To date, no normative explanation for such seemingly irrational behavior exists.

We propose that extinction burst is a result of the interplay between the learning and homeostatic systems. According to our theory, at the beginning of the extinction phase, the animal still expects to receive the outcome upon pressing the lever (Figure 3). During this period, due to the absence of an outcome, the internal state drops below the setpoint. This slight homeostatic deviation is sufficient to induce a transient increase in the response rate (i.e., burst), because the animal expects to receive the outcome and offset the internal deviation. Later on, as the animal learns that pressing the lever does not lead to the outcome, the rate of responding decreases in spite of aggravating homeostatic deviation.

**Figure 3.**
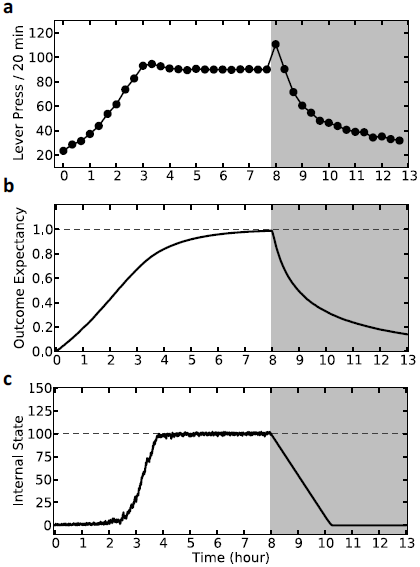
Simulation results demonstrating extinction burst. In every time step (every 4 s), the simulated agent can choose between pressing the lever (LP) or doing nothing (Figure S4). Pressing the lever results in an outcome with magnitude *k* = 5. Furthermore, in every time step, the internal state declines marginally, supposedly due to normal metabolism. The initial increase in response rate (a) is due to learning that pressing the lever results in an outcome (b). After the internal state reaches the setpoint, a stable response rate is maintained to preserve homeostasis (c). After 8 h, the outcome is removed, resulting in extinction burst followed by gradual suppression of responding.

In this respect, our model predicts that not only extinction, but even a reduction in the magnitude of the outcome will result in a temporary burst of responding (Figure S4). This prediction is in contrast to the classical homeostatic regulation models (see Materials and methods).

### Rescuing Hull?

Critically, our model is inspired by the drive reduction theory of motivation, initially proposed by Clark Hull (Hull, 1943), which became the dominant theory of motivation in psychology during the 1940s and 1950s. However, major criticisms have been leveled against this theory more recently (Berridge, 2004; McFarland, 1969; Savage, 2000). Our formal theory alleviates these major faults. In the earlier sections, we demonstrated that our theory redresses the first major fault of the classical drive-reduction: its inability to explain anticipatory responding in which animals paradoxically voluntarily increase (rather than decrease) their drive deviation, even in the absence of any physiological deficit. We demonstrated how such apparently maladaptive responses are learned and result in optimal behavior, ensuring physiological stability.

Second, the drive reduction could not explain how secondary reinforcers (e.g., money, or a light that predicts food) gain motivational value, since they do not reduce the drive *per se*. Because our framework integrates an RL module with the HR reward computation, the drive reduction-induced reward of primary reinforcers can be readily transferred through the learning process to secondary reinforcers that predict them (i.e., Pavlovian conditioning) as well as to behavioral policies that lead to them (i.e., instrumental conditioning).

Similarly, our integrated theory is able to give a normative account for the motivational effect of the orosensory components associated with primary physiological outcomes. To do so, we posit that sensory properties of food and water provide the animal with an unbiased estimate, *K̂_t_*, of their true post-ingestive effect, *K_t_*, on the internal state. Such association between sensory and post-ingestive properties could have been developed through learning or evolutionary mechanisms (Breslin, 2013).

Based on this sensory approximation, the reinforcing value of food and water outcomes can be approximated as soon as they are sensed/consumed, without having to wait for the outcome to be digested and the drive to reduce. In other words, the only information required to compute the reward (and thus the reward prediction error) is the current physiological state (*H_t_*) and the sensory-based approximation of the nutritional content of the outcome (*K̂_t_*):

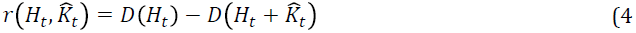

This proposition is in accordance with the fact that dopamine neurons exhibit instantaneous, rather than delayed, burst activity in response to unexpected food rewards (Schneider, 1989). Clearly, the evolution of the internal state itself (*H_t_* → *H_t_* + *K_t_*) depends only on the actual (*K_t_*) post-ingestive effects of the outcome.

This plausible sensory-estimate extension alleviates the other faults of drive reduction theory. Notably it explains the experimental fact that intravenous injection (and even intragastric intubation, in some cases) of food is not rewarding even though its drive reduction effect is equal to when it is ingested orally (Miller & Kessen, 1952) (*see also* (Ren et al., 2010)): the post-ingestive effect of food is estimated by its sensory properties and thus, the reinforcing value of intravenously injected food that lacks sensory aspects is zero. In the same line, the theory explains that animals’ motivation toward palatable foods, such as saccharine, that have no caloric content (and thus no need-reduction effect) is due to erroneous over-estimation of their drive-reduction capacity, misguided by their taste or smell.

A seminal series of experiments (McFarland, 1969) demonstrated that the reinforcing and satiating (i.e., need reduction) effects of drinking behavior, dissociable from one another, are governed by the orosensory and alimentary components of the water, respectively. Thirsty animals learned to peck at a key only when it resulted in oral, but bot intragastric (through a fistula) delivery of water (Figure 4 - figure supplement 1). Also, the response rate in the oral group initially increased but then gradually extinguished (rise-fall pattern; Figure 4 - figure supplement 1a). Simulating our theory (Figure 4) to account for this behavioral pattern (Figure 4a) clarifies that the ascending limb of the response curve represents a learning effect (Figure 4.c), whereas the descending limb is due to the internal state approaching the setpoint (i.e., satiation, as opposed “unlearning”; Figure 4e). Notably, classical RL models only explain the rise, and classical HR models only explain the fall. Furthermore, the simulation results show that the sensory component is crucial for approximating the drive-reduction reinforcing effect of water (Figure 4a,b). As above, the sensory-based approximation (*K̂_t_*) of the alimentary effect of water in the oral and fistula cases is assumed to be equal to its actual effect (*K_t_*) and zero, respectively (See Figure 4 - figure supplement 3 and 4, and Table S2 for simulation details).

**Figure 4.**
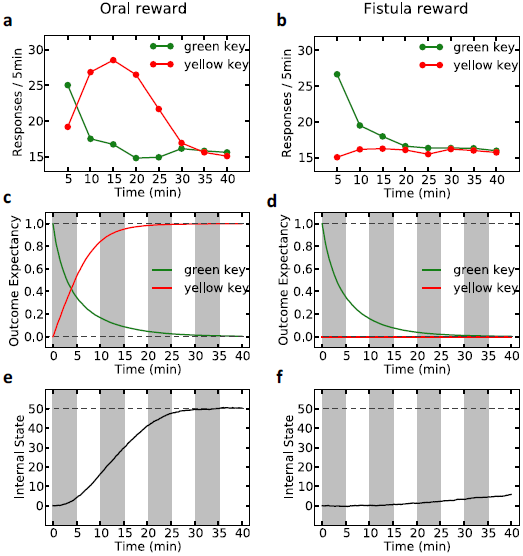
Simulation results replicating the data from (McFarland, 1969) on learning the reinforcing effect of oral vs. intragastric delivery of water. In a pre-training phase, simulated agents were trained in a task where responding on the green key resulted in the oral delivery of water. In the test phase, the green key had no consequence, whereas pecking at a novel yellow key resulted in oral delivery of water in one group (a) and intragastric injection of the same amount of water through a fistula in a second group (b). All agents started this phase in a thirsty state (initial internal state = 0; setpoint = 50). In the oral group, responding transferred rapidly from the green to the yellow key and was then suppressed (a) as the internal state approached the setpoint (e). This transfer is due to gradually updating the subjective probability of receiving water outcome upon responding on either key (c). In the fistula group, as the water is not sensed, the outcome expectation converges to zero for both keys (d) and thus, responding is extinguished (b). As a result, the internal state changes only slightly (f).

Our theory also explains why satiation is independent of the sensory aspects of water and only depends on its post-ingestive effects. Experiments show that when different proportions of water were delivered via the two routes in different groups, satiation (i.e., suppression of pecking) only depended on the total amount of water ingested, regardless of the delivery route (McFarland, 1969). Our model accounts for these data (Figure 4 - figure supplement 2), since evolution of the internal state only depends on the actual water ingested. Therefore, reaching the setpoint (and thus suppression of pecking) does not depend on the oral/fistula proportion. We predict, however, that the proportion affects the speed of satiation: higher oral proportions result in higher rewarding values of pecking, which in turn results in faster responding and thus faster satiation (Figure 4 - figure supplement 2).

### Behavioral plausibility of drive

The definition of the drive function (Eq. 1) in our model has two degrees of freedom: *m* and *n* are free parameters whose values determine the properties of the homeostatic space metric. Appropriate choice of *m* and *n* (*n* > *m* > 2) permits our theory to account for the following four key behavioral phenomena in a unified framework (see Materials and methods for mathematical derivations): (a) the potentiation of the reinforcing value of an appetitive outcome as its dose (*K_t_*) increases (Figure S5); (b) the excitatory effect of the deprivation level on reinforcing value (i.e., food will be more rewarding when the animal is hungrier; Figure S6); (c) the inhibitory effect of irrelevant drives (Figure S7), which is consistent with a large body of behavioral experiments showing competition between different motivational systems so that as the deprivation level for one need increases, it inhibits the rewarding value of other outcomes that satisfy irrelevant motivational systems (e.g., calcium deprivation reduces the appetite for phosphorus and hunger inhibits sexual behavior, etc.; see (Dickinson & Balleine, 2002) for a review); (d) finally, the theory naturally captures the risk-aversive nature of behavior. The rewarding value in our model is a concave function of the corresponding outcome amplitude (the second derivative of the reward with respect to the dose of outcome is negative). It is well known that the concavity of the utility function is equivalent to risk aversion (Mas-Colell, Whinston, & Green, 1995). Indeed, the model shows (Figure S8) that when faced with two options with equal expected payoffs, the model learns to choose the more certain option as opposed to the risky one. In fact, as an evolutionary function of risk-avoidance behavior, our theory provides the intuition that when the expected physiological instability of two behavioral options are equal, organisms do not choose the risky option, because the severe, though unlikely, physiological instabilities that it can cause might be life-threatening.

Our unified explanation for the above behavioral patterns implies that they all arise from the functional form of the mapping from the physiological to the motivational state.

### Neural substrates

Homeostatic regulation critically depends on sensing the internal state. In the case of energy regulation, for example, the arcuate nucleus of the hypothalamus integrates peripheral hormones including leptin, insulin, and ghrelin, whose circulating levels reflect the internal abundance of fat, abundance of carbohydrate, and hunger, respectively (Williams & Elmquist, 2012). In our model, the deprivation level has an excitatory effect on the rewarding value of outcomes and thus on the reward prediction error (RPE). Consistently, recent evidence indicates neuronal pathways through which energy state-monitoring peptides modulate the activity of midbrain dopamine neurons, which supposedly carry the RPE signal (Palmiter, 2007).

Namely, orexin neurons, which project from the lateral hypothalamus area to several brain regions, including the ventral tegmental area (VTA) (Sakurai et al., 1998) have been shown to have an excitatory effect on dopaminergic activity (Korotkova, Sergeeva, Eriksson, Haas, & Brown, 2003; Narita et al., 2006). Orexin neurons are responsive to peripheral metabolic signals as well as to the animal’s deprivation level (Burdakov, Gerasimenko, & Verkhratsky, 2005), as they are innervated by orexigenic and anorexigenic neural populations in the arcuate nucleus where circulating peptides are sensed. Accordingly, orexin neurons are suggested to act as an interface between internal states and the reward learning circuit (Palmiter, 2007). In parallel with the orexinergic pathway, ghrelin, leptin and insulin receptors are also expressed on the VTA dopamine neurons, providing a further direct interface between the HR and RL systems. Consistently, whereas leptin and insulin inhibit dopamine activity, ghrelin has an excitatory effect (see (Palmiter, 2007) for a review).

The reinforcing value of food outcome (and thus the RPE signal) in our theory is not only modulated by the internal state, but also by the orosensory information that approximates the need-reduction effects. In this respect, endogenous opioids and *μ*-opioid receptors have long been implicated in the hedonic aspects of food, signaled by its orosensory properties. Systemic administration of opioid antagonists decreases subjective pleasantness rating and affective responses for palatable foods in humans (Yeomans & Wright, 1991) and rats (Doyle, Berridge, & Gosnell, 1993), respectively. Supposedly through modulating palatability, opioids also control food intake (Sanger & McCarthy, 1980) as well as instrumental food-seeking behavior (Cleary, Weldon, O’Hare, Billington, & Levine, 1996). For example, opioid antagonists decrease the breakpoint in progressive ratio schedules of reinforcement with food (Barbano, Le Saux, & Cador, 2009), whereas opioid agonists produce the opposite effect (Solinas & Goldberg, 2005). This reflects the influence of orosensory information on the reinforcing effect of food. Consistent with our model, these influences have mainly been attributed to the effect of opiates on increasing extracellular dopamine levels in the Nucleus Accumbens (NAc) (Devine, Leone, & Wise, 1993) through its action on μ-opioid receptors in the VTA and NAc (Noel & Wise, 1993; M. Zhang & Kelley, 1997).

## Discussion

Theories of conditioning are founded on the argument that animals seek reward, while reward is defined as what animals seek. This apparently circular argument relies on the hypothetical and out-of-reach axiom of reward-maximization as the behavioral objective of animals. Physiological stability, however, is an observable fact. Here, we established a coherent mathematical theory where physiological stability is put as the basic axiom, and reward is defined in physiological terms. We demonstrated that reinforcement learning algorithms under such a definition of physiological reward lead to optimal policies that both maximize reward collection and minimize homeostatic needs. This argues for behavioral rationality of physiological integrity maintenance and further shows that temporal discounting of rewards is paramount for homeostatic maintenance. Furthermore, we demonstrated that such integration of the two systems provides normative explanations for several behavioral and neurobiological phenomena, including anticipatory responding, extinction burst, the rise-fall pattern of food-seeking response, and the modulation of midbrain dopaminergic activity by hypothalamic signals. We then extended the theory to incorporate orosensory information as an unbiased estimate of the post-ingestive effects of outcomes that is available instantaneously upon consumption. This extension of the theory allowed it to explain further behavioral patterns, rescue the classical drive-reduction theory, and shed light on the role of dopamine modulation by the opioid system.

From an evolutionary perspective, physiological stability and thus survival can themselves be seen as means of guaranteeing reproduction. These intermediate objectives can be even violated in specific conditions and be replaced with parental sacrifice. Still, homeostatic maintenance can explain a majority of motivated behaviors in animals. It is also noteworthy that our theory only applies to rewards that have a corresponding regulatory system. How to extend our theory to rewards without a corresponding homeostatic regulation system (e.g., social rewards, novelty-induced reward, etc.) remains a key challenge for the future.

Using internal state-independent reward/utility in the classical RL models and more generally in standard economic choice theory leads to the inevitable conclusion that maximizing utility is equivalent to maximizing the acquisition of appetitive outcomes. That is, more of a commodity equals more happiness. In opposition to this view, the rational decision in our model under certain circumstances is to choose small rather than large outcomes to avoid overshooting the setpoint (Figure S9). Furthermore, the theory explained several other key behavioral phenomena that stem from the interaction between the RL and HR systems: anticipatory responding, extinction burst, and the rise-fall pattern of responding.

Interestingly, a linear approximation of our proposed drive-reduction reward is equivalent to assuming that the rewarding value of outcomes is equal to the multiplication of the deprivation level and the magnitude of the outcome (see Materials and methods). In this respect, our model subsumes and provides a normative basis to the incentive salience theory (J. Zhang, Berridge, Tindell, Smith, & Aldridge, 2009) as well as other multiplicative forms of deprivation-modulated reward (e.g., decision field theory (Busemeyer, Townsend, & Stout, 2002), intrinsically motivated RL theory (Singh, Lewis, Barto, & Sorg, 2010), and MOTIVATOR theory (Dranias, Grossberg, & Bullock, 2008), where reward increases as a linear function of deprivation level.

Whether the brain uses a nonlinear drive-reduction reward (as in Eq. 2) or a linear approximation of it (as in Eq. 4) can be examined experimentally. Assuming that an animal is in a slightly deprived state (Figure 5a), a linear model predicts that as the magnitude of the outcome increases, its rewarding value will increase linearly. A non-linear reward, however, predicts an inverted U-shaped utility function (Figure 5b). That is, the rewarding value of a large outcome can be negative, if it results in overshooting the setpoint.

**Figure 5.**
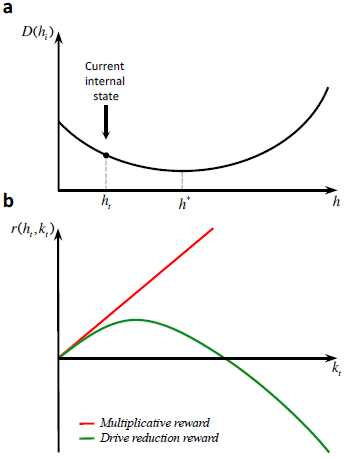
Differential behavioral predictions of multiplicative and drive-reduction forms of reward. Assuming that the internal state is at *h_t_* (a), outcomes larger than *h*\* – *h_t_* result in overshooting the setpoint and thus a declining trend of rewarding value (b).

In a nutshell, our theory incorporates a formal physiological definition of primary rewards into a novel homeostatically regulated reinforcement learning theory, allowing us to prove that economically rational behaviors ensure physiological integrity. At the same time our theory opens a unified approach behavioral pathologies (e.g., obesity, psychiatric diseases, drug addiction and compulsivity disorders) as allostatic dysregulation of the interactions between the motivational and learning brain axes.

## Acknowledgements

We thank Peter Dayan, Amir Dezfouli, Serge Ahmed, and Mathias Pessiglione for critical discussions and Oliver Hulme for commenting on the manuscript. This study was supported by fundings from Frontiers du Vivant, the French MESR, CNRS, INSERM, ANR, ENP and NERF. Support from the Basic Research Program of the Russian National Research University Higher School of Economics is gratefully acknowledged by BG.

---

**Figure 2 - figure supplement 1.**
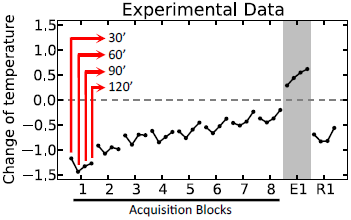
Experimental results (adapted from *(18)*) on the acquisition and extinction of conditioned tolerance response to ethanol. In each block (day) of the experiment, the animal receives ethanol injection after the presentation of the stimulus. The change in the body temperature is measured 30, 60, 90, and 120 minutes after ethanol administration. Initially, the hypothermic effect of ethanol decreases the body temperature of animals. After several training days, however, animals learn to activate a tolerance response upon observing the stimulus, resulting in smaller deviations from the temperature setpoint. If the stimulus is not followed by ethanol injection, as in the first day of extinction (E1), the activation of the conditioned tolerance response results in an increase in body temperature. The tolerance response gets weakened after several extinction sessions, resulting in increased deviation from the setpoint in the first day of re-acquisition (R1), where presentation of the cue is again followed by ethanol injection.

**Figure 2 - figure supplement 2.**
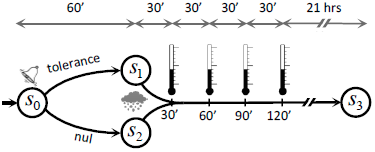
The model is simulated in an artificial task (Markov Decision Process) in which, upon observing the stimulus, the agent can choose between triggering the tolerance response and doing nothing.

**Figure 3 - figure supplement 1.**
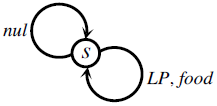
The Markov Decision Process used for simulating the extinction burst. At each time point, the agent can choose between doing nothing (*nul*) or pressing the lever (LP) which results in food delivery.

**Figure 4 - figure supplement 1.**
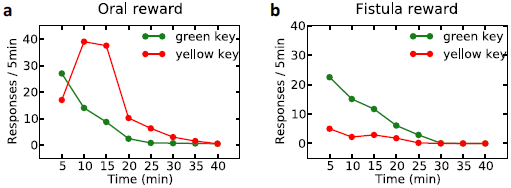
Experimental results (adapted from *(23)*) on learning the reinforcing effect of oral vs. intragastric delivery of water. Thirsty animals were initially trained to peck at a green key to receive water orally. In the next phase, pecking at the green key had no consequence, while pecking at a novel yellow key resulted in oral delivery of water in one group (a), and intragastric injection of the same amount of water through a fistula in a second group (b). In the first group, responding was rapidly transferred from the green to the yellow key, and then suppressed. In the fistula group, the yellow key was not reinforced.

**Figure 4 - figure supplement 2.**
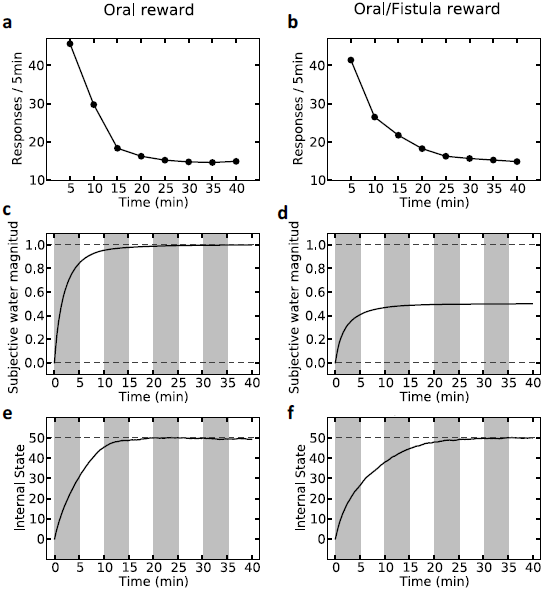
Simulation results of the satiation test. Left column shows results for the case where water was received only orally. Rate of responding drops rapidly (a) as the internal state approaches the setpoint (e). Also, the agent learns rapidly that upon every key pecking, it receives 1.0 unit of water (c). On the right column, upon every key-peck, 0.5 unit of water is received orally, and 0.5 unit is received via the fistula. As only oral delivery is sensed by the agent, the subjective outcome magnitude converges to 0.5 (d). As a result, the reinforcing value of key-pecking is less than the oral case and thus, the rate of responding is lower (b). This in turn results in slower convergence of the internal state to the setpoint (f).

**Figure 4 - figure supplement 3.**
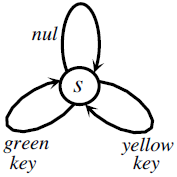
The Markov Decision Process used for simulating the reinforcing vs. satiation effects of water. At each time point, the agent can choose between doing nothing (*nul*) or pecking at either the green or the yellow key.

**Figure 4 - figure supplement 4.**
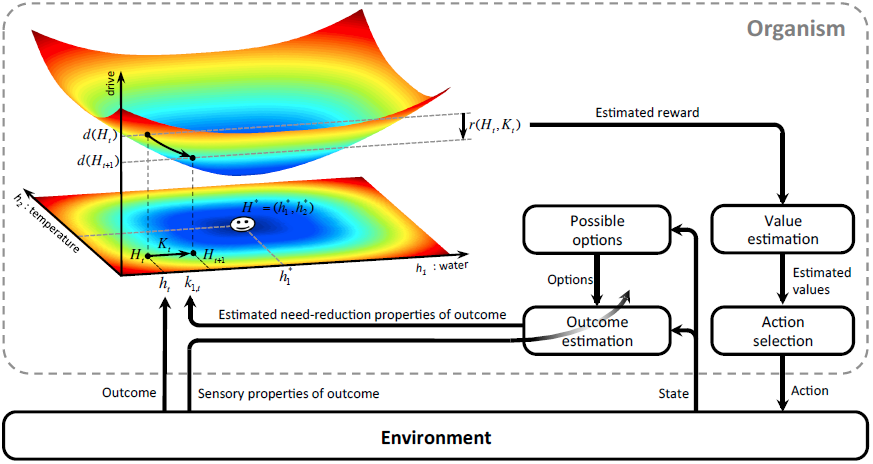
A model-based homeostatic RL system. Upon performing an action in a certain state, the agent receives an outcome, *K_t_*, which results in the internal state to shift from *H_t_* to *H_t_* + *K_t_*. At the same time, sensory properties of the outcome are sensed by the agent. Based on this information, the agent updates the state-action-outcome associations. In fact, the agent learns to predict the sensory properties, *K̂_t_*, of the outcome that is expected to be received upon performing a certain action. Having learned these associations, the agent can estimate the rewarding value of different options. That is, when the agent is in a certain state, it predicts the outcome *K̂_t_*, expected to result from each behavioral policy. Based on *K̂_t_* and the internal state *H_t_*, the agent can approximate the drive-reduction reward.

